# Precision medicine for the rescue of specific impairments in social behavior associated with the A350V *Iqsec2* mutation

**DOI:** 10.1101/2020.11.04.368282

**Authors:** Renad Jabarin, Nina Levy, Yasmin Abergel, Joshua H. Berman, Amir Zag, Shai Netser, Andrew P. Levy, Shlomo Wagner

## Abstract

In this study we tested the hypothesis that precision medicine guided therapy targeting glutamatergic neurotransmission could rescue behavioral deficits exhibited by mice carrying a specific mutation in the *Iqsec2* gene. The IQSEC2 protein plays a key role in glutamatergic synapses and mutations in the *IQSEC2* gene are a frequent cause of neurodevelopmental disorders. We have recently reported on the molecular pathophysiology of one such mutation A350V and demonstrated that this mutation downregulates AMPA type glutamatergic receptors (AMPAR) in A350V mice. Here we sought to identify behavioral deficits in A350V mice and hypothesized that we could rescue these deficits by PF-4778574, a positive AMPAR modulator. We found that A350V *Iqsec2* mice exhibit specific deficits in sex preference and emotional state preference behaviors as well as in vocalizations when encountering a female mouse. The social discrimination deficits, but not the impaired vocalization, were rescued with PF-4778574. We conclude that social behavior deficits associated with the A350V *Iqsec2* mutation may be rescued by enhancing AMPAR mediated synaptic transmission.

## Introduction

*IQSEC2* is an X-linked gene which has been previously associated with ASD, ID, and drug resistant epilepsy (1, 2). In this study we sought to demonstrate proof of concept for a precision medicine based program to rescue murine behavioral deficits associated with a specific missense mutation found in human *IQSEC2* (3) (replacement of alanine with valine at amino acid residue 350 of IQSEC2, denoted A350V *IQSEC2*).

The IQSEC2 protein contains a catalytic domain (Sec7) characteristic of all GEFS (guanine exchange factors), which promotes GDP exchange for GTP on Arf6. It also contains an IQ like domain, which has been proposed to bind calmodulin, and thereby modulate the Sec7 GEF activity of IQSEC2 (4–6). We have recently demonstrated that the A350V *IQSEC2* mutation results in the inappropriate constitutive activation of the Sec7 domain of IQSEC2, which leads to a marked downregulation of hippocampal surface AMPAR in A350V *Iqsec2* CRISPR generated mice (A350V mice) (7). We have further demonstrated that the decrease in surface AMPAR, specifically in the surface GluA2 receptor, is associated with a marked decrease in basal synaptic transmission in the murine model (7). These findings thereby identify AMPAR mediated transmission as a potential target for treating the deficits associated with the A350V mutation with precise medications. Therefore, we directly compared wild type (WT) and A350V mice in multiple different social discrimination paradigms (8, 9) as well as in several types of social vocalization. We then assessed the ability of a positive allosteric modulator (PAM) of AMPAR, which serves to increase AMPAR mediated synaptic transmission by slowing AMPAR desensitization (10, 11) to rescue impairments in these behaviors exhibited by A350V mice.

## Methods and Materials

### Study design

The initial objective of this study was to identify deficits in social behavior exhibited by A350V *Iqsec2* mice, as compared to their WT littermates. Following the accomplishment of this objective, we defined the assessment of the short- (1-2 days) and long-term (7-11 days) effect of PF-4778574 treatment, as compared to vehicle injection, on these deficits as another objective. The initial sample size for behavioral experiments was determined to be 20±5, based upon power calculations made in a previous study using C57BL/6J mice (9). For behavioral deficits in *Iqsec2* mice, we repeated the results in two independent cohorts, so overall sample size was double. The effect of vehicle treatment on SxP and ESP in both WT and A350V mice was examined at 1-2 days and 7-8 days following injection. The effect of PF-4778574 treatment on A350V mice was examined at 1-2 days (SxP + ESP) and again at 7-8 days (SxP) or at 11 days (ESP) following treatment. The results of PF-4778574 treatment was qualitatively replicated in two independent cohorts, with slightly smaller sample size due to animal availability. No animals were excluded from experiments and no outliers were defined. A350V and WT mice were always examined together on the same day in a random order. All data collection was fully automated, with no involvement of an observer.

### Animals

C57BL\6J subjects were commercially obtained (Envigo, Israel) naïve males (8-12 weeks old, 3-5/cage). Social stimuli were in-house grown C57BL\6J juvenile (21-30 days old males (SP, SNP) or naïve adult male and female mice (SxP, ESP). Emotionally aroused stimuli for the ESP paradigm were each individually housed for 1 week.

A350V *Iqsec2* mice generation, maintenance and genotyping were previously described (7). All *Iqsec2* animal were bred and maintained in the animal facility of the faculty of medicine of the Technion Institute of Technology until age of 8-10 weeks and then transferred to the University of Haifa mice facility for at least one week before experiments.

All animals were kept on a 12-h light\12-h dark cycle, light on at 7p.m., with *ad libitum* access to food and water. Behavioral experiments took place during the dark phase under dim red light. All animal protocols and experiments were approved by institutional animal care and use committees (IL0460416; IL1691117; IL1271118).

### PF-4778574 Preparation and Injection

PF-4778574 [N-<(3R,4S)-3-[4-(5-cyano-2-thienyl)phenyl]tetrahydro-2H-pyran-4-yl>propane-2-sulfonamide] was obtained as a powder from Sigma (catalog #PZ0211) and solubilized (2.5 mg/ml) in dimethyl sulfoxide (sigma cat # D2650). Aliquots were stored at −20°C and diluted immediately prior to injection in 5% dimethyl sulfoxide, 5% Kolliphor EL (Sigma cat #C5135) and 90% dH20 as previously described (11). The PF-4778574 (0.30 mg/kg) and vehicle solutions (200 ul) were administered subcutaneously and subject mice were returned to their home cage until the day of experiment.

### Behavioral assays

All social discrimination tasks were conducted using our published automated experimental system (8). SP and SNP tests were conducted on the same day, as previously described (9).

The SxP test consisted of 15 min habituation to the arena with empty chambers, followed by exposing the subject for 5 min to both adult male and female social stimuli located in individual chambers at opposite corners of the arena.

The ESP test similarly consisted of a 15 min habituation, followed by exposing the subject for 5 min to both group-housed and socially-isolated (1 wk.) stimuli confined to individual chambers randomly located at opposite corners of the arena. Each isolated stimulus was used for two nonconsecutive tests. All stimuli used in all four tests (SP, SNP, SxP and ESP) were C57BL\6J mice.

### Vocalizations recording

Ultrasonic vocalizations were recorded using a condenser ultrasound microphone (Polaroid/CMPA, Avisoft) placed above the experimental arena and sampled at 250 kHz. Vocalizations were recorded during a 10 min interaction with a female stimulus, following a 10 min of habituation to the arena. For the recording of pup calls, pups were moved to a netted metal cup above which the microphone was placed, vocalizations were recoded for a period of 2 min.

### Data analysis

Video data analysis was performed by our published custom-made TrackRodent software, as previously described (8). Audio data was analyzed using our TrackUSF custom-made software as described in https://www.biorxiv.org/content/10.1101/575191v1.

### Statistical analysis

SPSS v21.0 (IBM) was used for statistical analysis. Following a Kolmogorov–Smirnov test for checking normal distribution of the dependent variables, a 2-tailed paired t-test was used to compare between parameters within a group, and a 2-tailed independent t-test was used to compare a single parameter between distinct groups. Otherwise, the non-parametric Mann-Whitney U test was used to compare between parameters in the same group. For comparison between multiple groups and parameters a mixed analysis of variance (ANOVA) model was applied to the data. This model contains one random effect (ID), one within effect, one between effect, and the interaction between them. For comparison within a group using multiple parameters, a two-way repeated measures ANOVA model was applied to data. This model contains one random effect (ID), two within effects, one between effect and the interactions between them. All ANOVA tests were followed, if main effect or interaction were significant, by *post hoc* Student’s t test. Significance was set at 0.05.

## Results

### Social behavior of C57BL\6J adult male mice

For analyzing the social behavior of mice subjects in this study, we have used our previously described experimental system which allows automated and detailed analysis of discrimination behavior (8) across a battery of four social discrimination tests. First, we have used the social preference (SP) test to assess social motivation and the social novelty preference (SNP) test to assess social recognition. We have also used the sex preference (SxP) test, in which the subject is exposed simultaneously to male and female conspecifics. Finally, we have used a novel test, termed the emotional state preference (ESP) test, in which the subject is exposed simultaneously to socially-isolated and group-housed mice. Figure 1 shows the results of all these tests when conducted with C57BL/6J adult male mice. In the SP test (Fig. 1A) these mice showed a significant preference of the social stimulus over the object throughout the test (Fig. 1B-C; paired t-test - t_19_=4.475, p<0.001). In the SNP test (Fig. 1D), the same subject mice showed a significant preference towards the novel mouse over the familiar one (Fig. 1E-F; paired t-test - t_19_=-2.66, p<0.05). In the SxP test (Fig. 1G), C57BL\6J mice spent significantly more time investigating a female over a male stimulus (Fig. 1H-I; paired t-test- t_24_=5.137, p<0.001), indicating a preference for mice of the opposite sex. As for the ESP test (Fig. 1J), C57BL\6J subjects showed a preference to investigate the socially-isolated (“emotional”) mouse over the group-housed mouse (Fig. 1K-L; paired t-test - t_13_=3.03, p<0.01), suggesting a tendency to socialize more with mice which are emotionally aroused (12, 13). As previously reported by us (9), in all tasks the subject’s preference was expressed mainly by a strong bias in long (>6 sec) investigation bouts, while we observed no preference between the stimuli when short (≤6 sec) bouts were considered (Fig. 1M-P).

**Fig. 1.**
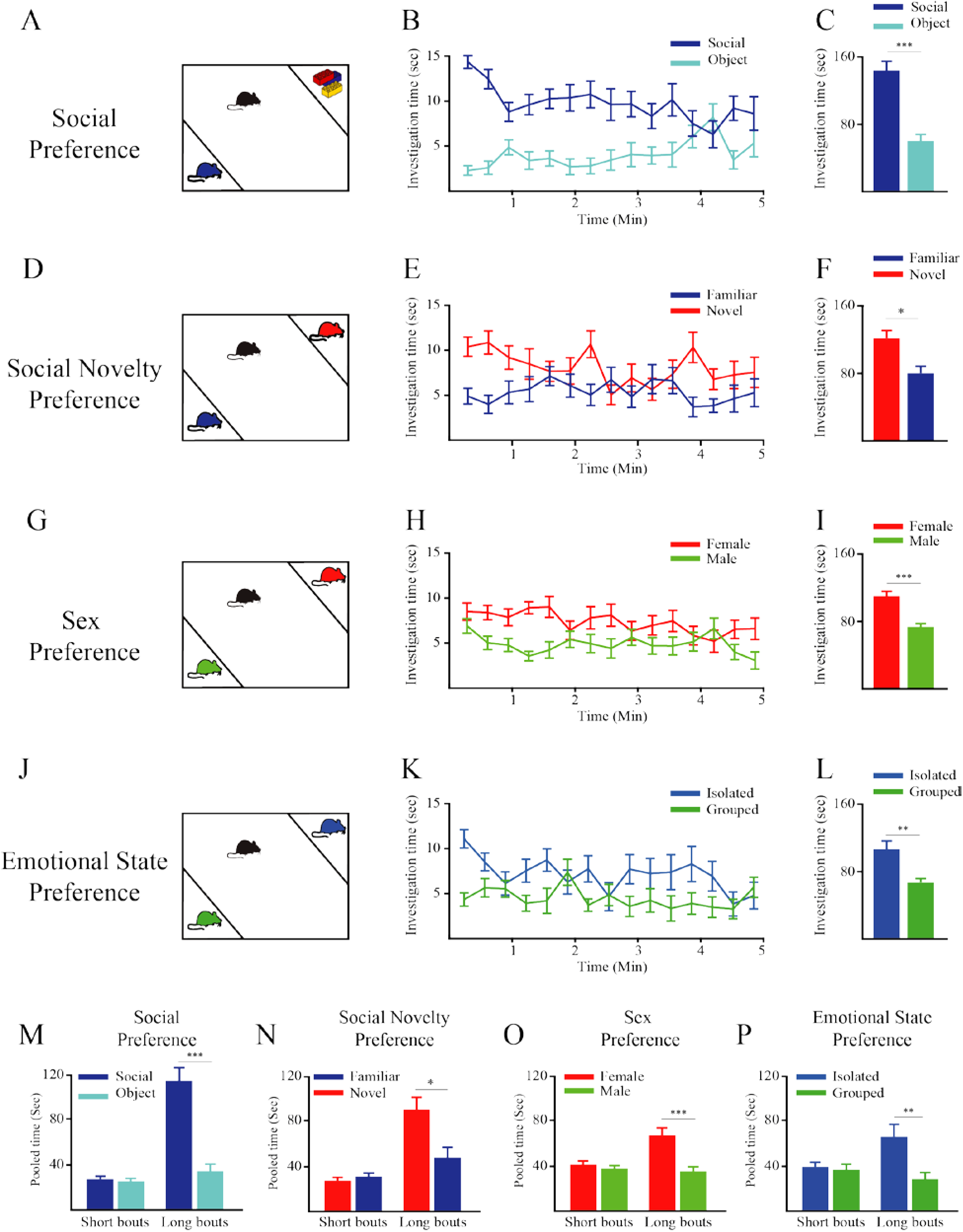
Social discrimination tests conducted with adult C57BL\6J male mice. (A) Schematic depiction of the social preference (SP) test. (B) Mean investigation time measured separately for each stimulus (20-s bins) across the SP test (n = 20). (C) Mean investigation time summed separately for each stimulus throughout the SP test. (D-F) As in A-C, respectively, for the social novelty preference (SNP; n=20) test. (G-I) As in A-C, respectively, for the sex preference (SxP; n=25) test. (J-L) As in A-C, respectively, for the emotional state preference (ESP; n=14) test. (M-P) Mean investigation time for each of the stimuli, pooled separately for short (<7 sec) and long (≥7sec) investigation bouts, for the SP (M), SxP (O) and ESP (P) tests shown above (A-L). Note that the preference for a specific stimulus is reflected only by the long bouts in all of these tests. *Post hoc* paired t-tests following main effect in mixed model ANOVA test; M-short bouts: t_19_=-0.576, p=0.571, long bouts: t_19_=-4.64, p<0.001; N-short bouts: t_19_=-0.93, p=0.364, long bouts: t_19_=2.78, p<0.05; O-short bouts: t_24_=0.973, p=0.34, long bouts: t_24_=4.825, p<0.001; P-short bouts: t_13_=0.708, p=0.492, long bouts: t_13_=3.167, p<0.01 *p<0.05, **p<0.01, ***p<0.001, paired t-test following main effect in ANOVA test.

### A350V Iqsec2 male mice exhibit specific deficits in social behavior

Next, we assessed the behavior of wild-type (WT) and A350V male littermates in the same battery of tests as described above for C57BL/6J mice. We found that both genotypes showed intact social preference in the SP test, with significantly greater investigation time for the social stimulus over the object (Fig. 2A-C, mixed-model ANOVA - within stimulus: F_(1,37)_=10.06, p<0.01; between genotypes: F_(1,37)_=0.259, p=0.614; stimulus×genotype: F_(1,37)_=0.142, p=0.709; *post hoc* paired t-test - WT: t_18_=2.178, p<0.05; A350V:t_19_=2.357, p<0.05). There was also no difference between the two genotypes in the amount of time spent investigating each of the stimuli (independent samples t-test - social: t_36.9_=0.557, p=0.581; object: t_37_=-0.007, p=0.994), suggesting that the A350V Iqsec2 mutation did not affect the motivation of the animals for social interactions. In the SNP test, both genotypes showed a similar preference for investigating the novel stimulus as observed in C57BL\6J mice (Fig. 2D-F; mixed-model ANOVA - within stimulus: F_(1,37)_=23.3, p<0.001; between genotypes: F_(1,37)_=0.002, p=0.966, stimulus ×genotype: F_(1,37)_=0.106, p=0.746; *post hoc* paired t-test - WT: t_18_=-3.721, p<0.01; A350V: t_19_=-3.13 p<0.01). Here too there was no difference between the WT and A350V littermates in the amount of time spent investigating each of the stimuli (t-test - Familiar: t_37_=-0.274, p=0.785, Novel: t_37_=0.275, p=0.785).

**Fig. 2.**
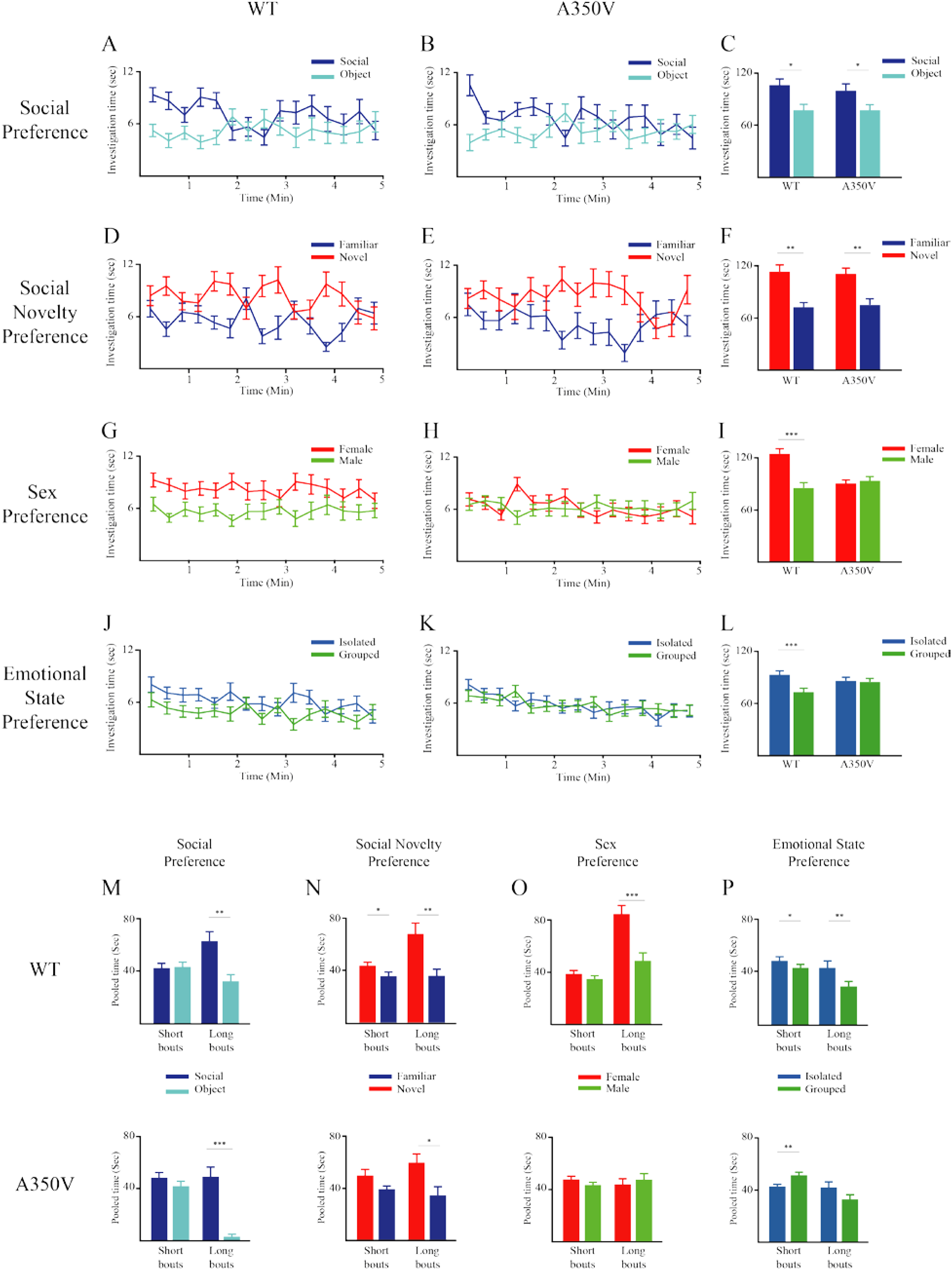
Deficits in social behavior of A350V *Iqsec2* mice. (A-B) Mean investigation time measured separately for each stimulus (20-s bins) across the SP test for WT (A, n=19) and A350V (B, n=20) mice. (C) Mean investigation time summed separately for each stimulus throughout the SP test for WT (left) and A350V (right) subjects. (E-F) Same in A-C, for the SNP test (WT - n=19, A350V – n=20). (G-I) Same in A-C, for the SxP test (WT - n=33, A350V – n=36). (J-L) Same in A-C, for the ESP test (WT - n=28, A350V – n=43). (M-P) Mean investigation time of WT (above) and A350V (below) mice for each of the stimuli, pooled separately for short (<7 sec) and long (≥7sec) investigation bouts, for the SP (M), SNP (N), SxP (O) and ESP (P) tests shown above (A-L). Note that A350V mice showed an atypical significant preference for the group-housed stimulus when short bouts were considered. *Post hoc* paired t-tests following main effect in mixed model ANOVA test; M-WT: short bouts: t_18_= −0.159, p=0.876, long bouts: t_18_=3.109, p<0.01; A350V: short bouts: t_19_=1.277, p=0.217, long bouts: t_19_=6.7, p<0.001; N-WT: short bouts: t_18_=-2.286, p<0.05, long bouts: t_18_=-2.897, p<0.01; A350V: short bouts: t_19_=-1.954, p=0.066, long bouts: t_19_=-2.4, p<0.05; O-WT: short bouts: t_32_= 1.157, p=0.256, long bouts: t_32_=3.65, p<0.001; A350V: short bouts: t_35_= −0.278, p=0.776, long bouts: t_35_=-0.243, p=0.809; P-WT: short bouts: t_27_= 2.11, p<0.05, long bouts: t_27_=3.015, p<0.01; A350V: short bouts: t_42_= −3.23, p<0.01, long bouts: t_42_=1.76, p=0.086. *p<0.05, **p<0.01, ***p<0.001, paired t-test following main effect in ANOVA test.

In contrast, we found a significant deficit displayed by A350V mice compared to WT littermates in the SxP test. While WT male mice, similarly to C57BL\6J male mice, showed a clear preference for females, A350V male mice showed no preference for either sex (Fig. 2G-I; mixed-model ANOVA - stimulus×genotype: F_(1,67)_=11.18, p<0.001; *post hoc* paired-samples t-tests - WT: t_33_=3.895, p<0.001; A350V: t_35_=-.416, p=0.68). There was no difference between the two genotypes when the total investigation time of both stimuli was considered (Mann-Whitney test - z=-0.207, p=0.836), indicating that the deficit in sex preference behavior observed in A350V mice is not due to an overall decrease in social motivation. The difference in sex preference behavior between WT and A350V littermates was consistent across two cohorts of subjects (Fig. S1A-F).

A similar difference between WT and A350V littermates was observed in the ESP test. While WT mice showed a significant preference for investigating the isolated stimulus, similarly to C57BL\6J mice, A350V mice showed no preference for either stimuli (Fig. 2J-L; mixed-model ANOVA - stimulus×genotype: F_(1,69)_=4.329, p<0.05; *post hoc* paired t-test - WT: t_27_=4.904, p<0.001; MT: t_42_=0.171, p=0.865). Here too, there was no difference between WT and A350V mice in the total investigation time of both stimuli (Mann-Whitney test - z=-0.353, p=0.724). This difference in ESP between WT and A350V mice was consistent across two cohorts of subjects (Fig. S1G-L). Interestingly, when analyzing the data separately for short and long bouts (Fig. 2M-P) we found that A350V mice showed an atypical significant preference for the grouped housed stimulus (Fig. 2P-lower; 2-way ANOVA-stimulus ×bout duration: F_(1,42)_=12.014, p<0.001; *post hoc* paired t-test-short bouts: t_42_=-3.23, p<0.01, long bouts-t_24_=1.76, p=0.086).

We conclude that while A350V *Iqsec2* mice behave the same as their WT littermates in the SP and SNP tests, they exhibit specific deficits in the SxP and ESP tests.

### A350V Iqsec2 male mice exhibit a specific deficit in mating calls

Male mice are known to emit ultrasonic vocalizations (USVs) when introduced to female mice (mating calls), but not to male mice (14). We recorded mating calls of WT and A350V littermates and analyzed them using our novel analysis system (https://www.biorxiv.org/content/10.1101/575191v1) which enables separation of ultrasonic audio fragments (USFs) from noise signals and their clustering using a three-dimensional t-SNE analysis. We found that while a large number of WT animals emitted a reasonable amount of ultrasonic vocalizations, none of their A350V littermates exhibited such behavior (Fig. 3A-C) and this difference was statistically significant (Fig. 3D; Mann-Whitney test-U=399, p<0.05). In contrast, there was no apparent difference in the noise generated by the WT and A350V littermates (Fig. 3A), suggesting no distinction in viability and movement between the two groups. In order to demonstrate that these results were reproducible, we conducted a similar experiment with a new cohort of animals in a different laboratory by distinct research staff and obtained similar results (Fig. S2; Mann-Whitney test-U=39, p<0.05). In order to verify that A350V mice are capable of emitting ultrasonic vocalizations, we recorded pup separation calls from WT and MT littermates and found no significant difference between them (Fig. S3; Kruskal Wallis test-chi=0.469, p=0.791). Altogether, these results suggest a specific deficit exhibited by A350V *Iqsec2* male mice in the emission of mating calls during interaction with a female mouse.

**Figure 3.**
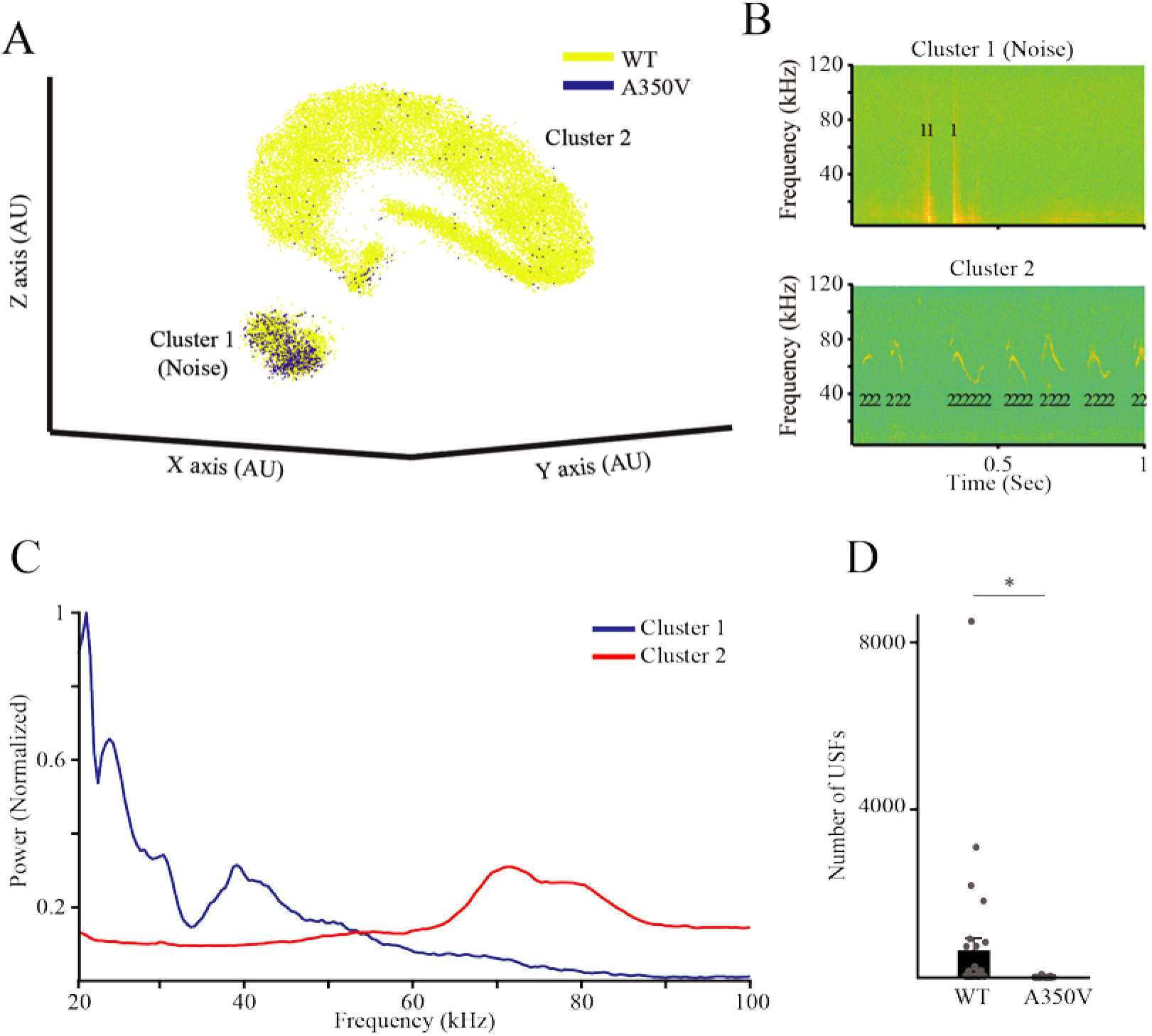
A350V *Iqsec2* male mice do not emit mating calls. (A) 3D t-SNE analysis of all ultrasonic fragments emitted by WT (yellow; n=31) and A350V (blue; n=36) adult male mice during an encounter with a female C57BL/6J mouse. Note that while cluster 1 (noise) seem to contain similar number of fragments from both genotypes, cluster 2 (vocalizations) contains almost only fragments from WT mice. (B) Examples of USFs, denoted by their cluster numbers, superimposed on the spectrograms of their respective audio signals. Note that USFs of cluster 1 (upper example) represent non-vocal (noise) signals while USFs of cluster 2 (lower example) represent genuine vocalizations. (C) Power Spectral Density (PSD) analysis of all USFs recorded from all animals, calculated separately for each cluster. Note that in contrast to the PSD profile of cluster 1 which shows a wide range of frequencies, mainly at the lower range, the profile of cluster 2 shows a clear peak at the range of 60-90 kHz. (D) Comparison of the number of USFs from cluster 2 between the two genotypes. Note that A350V mice emitted significantly lower number of USFs as compared to WT animals (Mann-Whitney U test. U=399.000, *p<0.05).

### Rescuing social behavior deficits with PF-4778574

We previously revealed a specific deficit in surface expression of the GluA2 AMPAR in the hippocampus of A350V *Iqsec2* mice, which was accompanied by a reduction in glutamatergic synaptic activity in this region (7). Since the AMPAR modulator PF-4778574 was able to rescue social behavioral deficits in *Cntnap2*-knockout mice, which also display depressed glutamatergic transmission (15), we reasoned that enhancing AMPAR activity with PF-4778574 might alleviate the behavioral impairments exhibited by A350V mice. To examine this hypothesis, we assessed the behavior of A350V mice following vehicle administration as well as 1 day and 1-2 weeks following PF-4778574 administration.

In the SxP test, WT mice showed intact sex preference 1 or 2 days following vehicle injection (Fig. 4A; t_13_=2.163, p<0.05, paired t-test). In contrast, no such preference was observed among either vehicle or drug injected A350V mice 1 or 2 days following vehicle injection (Fig. 4B; mixed-model ANOVA - within stimulus: F_(1,50)_=0.051, p=0.821; between conditions: F_(1,50_)=0.084, p=0.773, stimulus×condition: F_(1,50)_=0.353, p=0.550). Nevertheless, 7 or 8 days after treatment drug injected A350V mice exhibited normal sex preference, in contrast to vehicle injected animals (Fig. 4C; mixed-model ANOVA - within stimulus: F_(1,36)_=8.382, p=0.006; between conditions: F_(1,36)_=0.471, p=0.497, stimulus×condition: F_(1,36)_=1.282, p=0.265; *post hoc* paired t-test: vehicle: t_11_=1.974, p=0.074; drug: t_25_=3.129, p<0.01).

**Figure 4.**
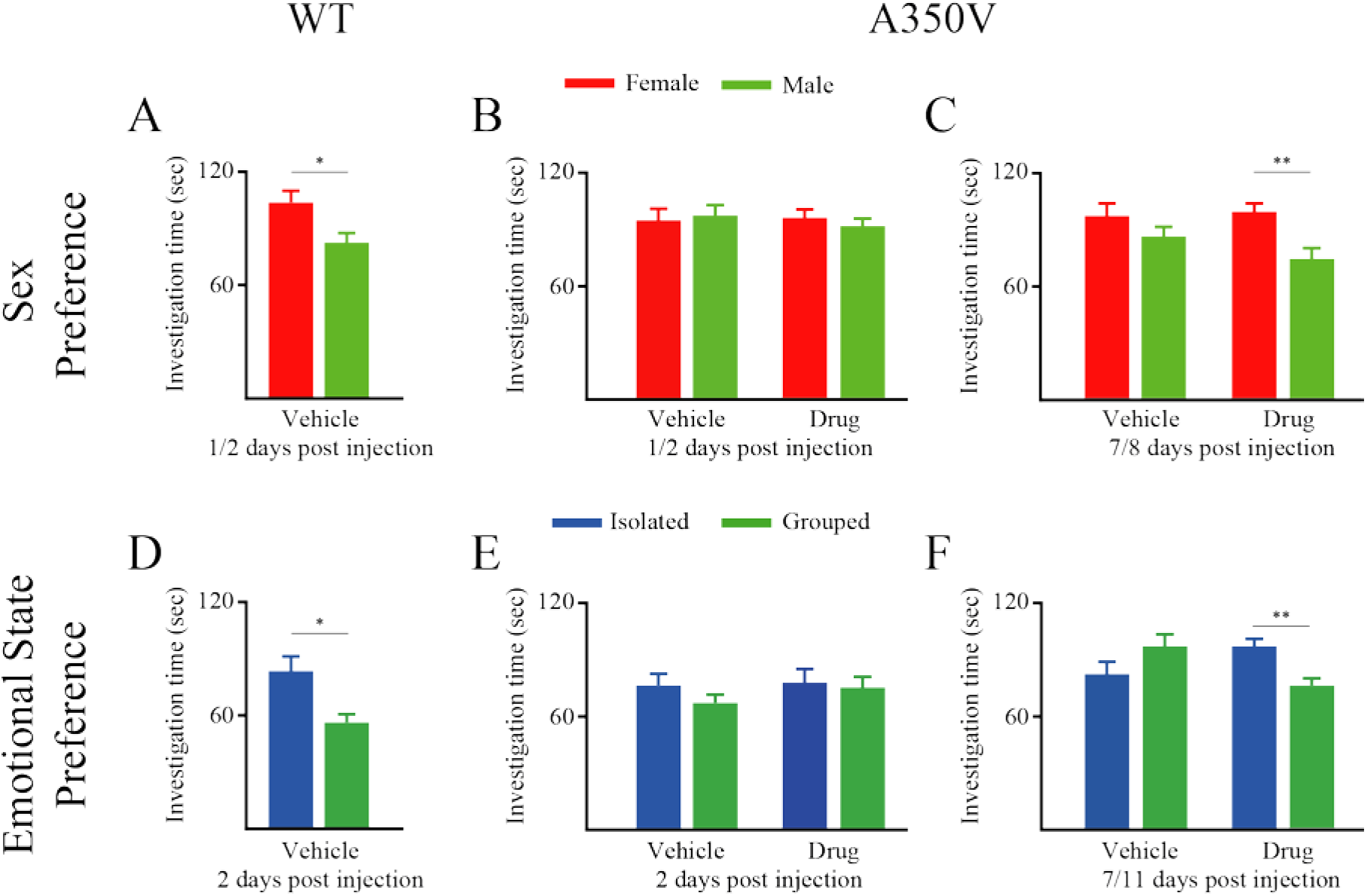
PF-4778574 rescues sex social behavioral deficits in A350V *Iqsec2* mice. (A) Mean investigation time summed separately for the Female and Male stimuli throughout the SxP test, conducted by WT mice 2 days after vehicle injection. (B) Mean investigation time summed separately for the Female and Male stimuli throughout the SxP test conducted by A350V mice 1-2 days after either vehicle (left, n=26) or PF-4778574 (right, n=26) injection. (C) As in B, for the SxP test conducted by A350V mice 7-8 days after either vehicle (left, n=12) or PF-4778574 (right, n=26) injection. (D) Mean investigation time summed separately for the Isolated and Grouped stimuli in the ESP test conducted by WT mice 2 days after vehicle injection. (E) Mean investigation time summed separately for the Isolated and Grouped stimuli in the ESP test conducted by A350V mice 1-2 days after either vehicle (left, n=12) or PF-4778574 (right, n=26) injection. (F) As in E, for the ESP test conducted by A350V mice 7-11 days after either vehicle (left, n=12) or PF-4778574 (right, n=26) injection. *p<0.05, **p<0.01, paired t-test following main effect in ANOVA test.

A similar benefit of PF-4778574 in A350V mice was obtained in the ESP test. WT mice showed clear preference towards the isolated social stimulus 1 or 2 days following vehicle injection (Fig. 4D; t_15_=2.65, p<0.05, paired t-test). In contrast, neither vehicle nor drug injected A350V mice showed preference between the two stimuli when tested 1 or 2 days following treatment (Fig. 4E; mixed-model ANOVA - within stimulus: F_(1,36)_=0.01, p=0.922; between conditions: F_(1,36)_=1.582, p=0.217; stimulus×condition: F_(1,36)_=0.04, p=0.842). However, when tested 7 or 11 days after treatment, drug injected A350V mice exhibited a significant preference for the isolated stimulus, in contrast to vehicle injected A350V mice (Fig. 4F; mixed-model ANOVA - stimulus×condition: F_(1,36)_=6.813, p=0.013; *post hoc* paired t-test - vehicle: t_11_=--1.28, p=0.227, drug: t_25_=2.766, p<0.01).

Finally, we examined the effect of PF-4778574 administration on the lack of mating calls observed in A350V mice. We found no improvement in social vocalizations 1 hour and 7 days following drug administration (Fig. S4).

## Discussion

In this study we have employed a battery of social tests to characterize the deficits in social behavior exhibited by mice with a A350V mutation in the *Iqsec2* gene, a mutation which is associated with epilepsy, ID and ASD in humans. We also demonstrated that a single administration of PF-4778574, a PAM modulator of the AMPAR, can rescue some of these deficits.

Numerous missense and nonsense mutations have been identified in the X-kinked *IQSEC2* gene which have been associated with ID, ASD and drug resistant epilepsy (1). Specific genotype-phenotype correlations have not been identified to date. Ultimately for the purposes of precision medicine treatment of *IQSEC2* mutations it will be necessary to understand if the molecular pathophysiology of these mutations differs. The majority of human mutations are nonsense mutations and several recent studies in which IQSEC2 expression was manipulated in neurons (5, 16) or the gene knocked out in mice models (17, 18) may be informative in understanding the molecular pathophysiology and phenotypes of nonsense mutations as compared to the A350V *IQSEC2* missense mutation (7). In vitro studies in neurons in which the catalytic activity of the Sec7 domain was inhibited or constitutively activated suggested that inhibition of the Sec7 activity of IQSEC2 would lead to an impairment in Arf6 mediated AMPAR recycling and consequently increased surface AMPAR expression (5, 16) while mutations which appeared to increase Arf6 activation would result in downregulation of AMPAR expression as observed in A350V *Iqsec2* mice (7). The phenotype of mouse knockout models of *Iqsec2* (17, 18) appears to have several notable differences from the A350V *Iqsec2* mutation phenotype as investigated in this and prior studies. First, regarding epilepsy, both knockout models have reported seizures occurring at approximately day 90 while with the A350V mutation seizures are limited to days 14-21 (4). Second, both knockout models have reported an increase in anxiety related behavior which we have not observed in A350V mice (7). Differences between the knockout model and the A350V model are not surprising given that the A350V mutation appears to enhance the Sec7 activity of IQSEC2. Demonstration of such differences between loss of function and gain of function in IQSEC2 implies that treatments that may be beneficial for one mutation of *IQSEC2* may not be effective (or even harmful) for a different mutation in this gene. Indeed such a dichotomy has been demonstrated for another gene controlling AMPAR regulation in which a knockout mutation in the *Thorase* gene resulted in an upregulation of surface AMPAR expression while a gain of function mutation in the same gene resulted in a downregulation in surface AMPAR (19–21).

In a previous study using the same mouse model used here, we demonstrated a normal sociability and social novelty preference in the three-chamber test (7). Here, we employed a novel experimental system designed for directly monitoring social investigation behavior to examine the behavior of these mice in a battery of four social discrimination tests. Of these tests, the ESP test is a novel test, which is similar in principle to the affective state discrimination test recently described (12), and seems to be highly relevant to ASD symptoms which involve impaired theory of mind (22, 23). In agreement with our previous paper, we found normal behavior of A350V mice in the SP test which assesses social motivation, as well as in the SNP test which assesses social cognition. We did find, however, that A350V mice did not discriminate between the stimuli in both the SxP and ESP tests, suggesting an impairment in the behavior of A350V mice towards social stimuli which are emotionally attractive, such as opposite sex and aroused stimuli. Future studies will find whether this is due to an impaired recognition of such stimuli or the lack of motivation to interact with them.

Our observation that A350V *Iqsec2* mice had a dramatic reduction mating calls during male-female interactions, as opposed to their WT littermates, is in agreement with a previous study characterizing *Iqsec2*-knockout mice (18). Such a strong deficit in mating calls was previously described for several genetic mouse models of ASD (24, 25). Nevertheless, PF-4778574 administration did not have any effect on this impairment, despite the rescue of sex preference behavior. This suggests that the lack of mating calls in A350V *Iqsec2* male mice is not due to their lack of motivation for social interactions with females, demonstrated by their impaired SxP behavior, but rather is likely due to a deficit in their social communication.

Benefits from PF-4778574 in the SxP and ESP tests were not achieved immediately (1-2 days post administration) but rather 7-11 days following administration. In this point our results differ from those previously described for *Cntnap2*-knockout mice, where the behavioral effects were observed already 30 min post drug administration (15). Future studies are needed in order to reveal what is the mechanism by which PF-4778574 induces its delayed and long lasting effect. Moreover, our study was limited to a single protocol of PF-4778574 application, with the drug’s dose and route of administration selected based on previous studies demonstrating its efficacy in rescuing AMPAR mediated memory impairments in wild type animals treated with ketamine (11) and social behavioral defects in *Cntnap2*-knockout mice (15). Thus, future studies are needed for the development of an optimal administration protocol that will allow maximal benefits from this drug or similar modulators of AMPAR activity.

To conclude, we have utilized a battery of social discrimination tasks to analyze modified social behavior in A350V Iqsec2 mutant mice. We observed specific deficits in the social behavior of these mice, which suggest that these mice are impaired in socio-emotional behavior and social communication. Moreover, we found that a single administration of the AMAP receptor PAM PF-4778574 rescued the behavioral deficits, while having no effect on the deficits in social vocalization exhibited by A350V male mice. Decreased AMPAR mediated synaptic transmission appears to represent a convergent pathway for an increasing number of genetic syndromes, including Fragile X (26), Rett (27), Cdkl5 (28), Rab39B (29), Shank (30), tuberous sclerosis (31), Atad1 (19–21), and Cntnap2 (15) as well as in non-genetic syndromes such as neonatal hypoxia (32), suggesting that it may be possible to extrapolate treatments from one of these disorders to the others. The findings of this study may therefore be broadly relevant for a large number of neurodevelopmental disorders of different etiologies and not just for this one mutation in *IQSEC2*.

## Acknowledgments

This study was supported by The Human Frontier Science Program (HFSP grant RGP0019/2015 for SW), the Israel Science Foundation (ISF grants #1350/12, 1361/17 for SW), by a donation by the Milgrom family for SW and by the Ministry of Science, Technology and Space of Israel (Grant #3-12068 for SW).

## Disclosures

### Author contributions

Study conception and design – SW, APL; Acquisition of data – RJ, NL, YA, JHB, AZ; Analysis and interpretation of data – RJ, SN; Drafting of manuscript – SW, APL.

### Competing interests

The authors declare no competing interest.

## Supplementary Figures

**Figure S1.**
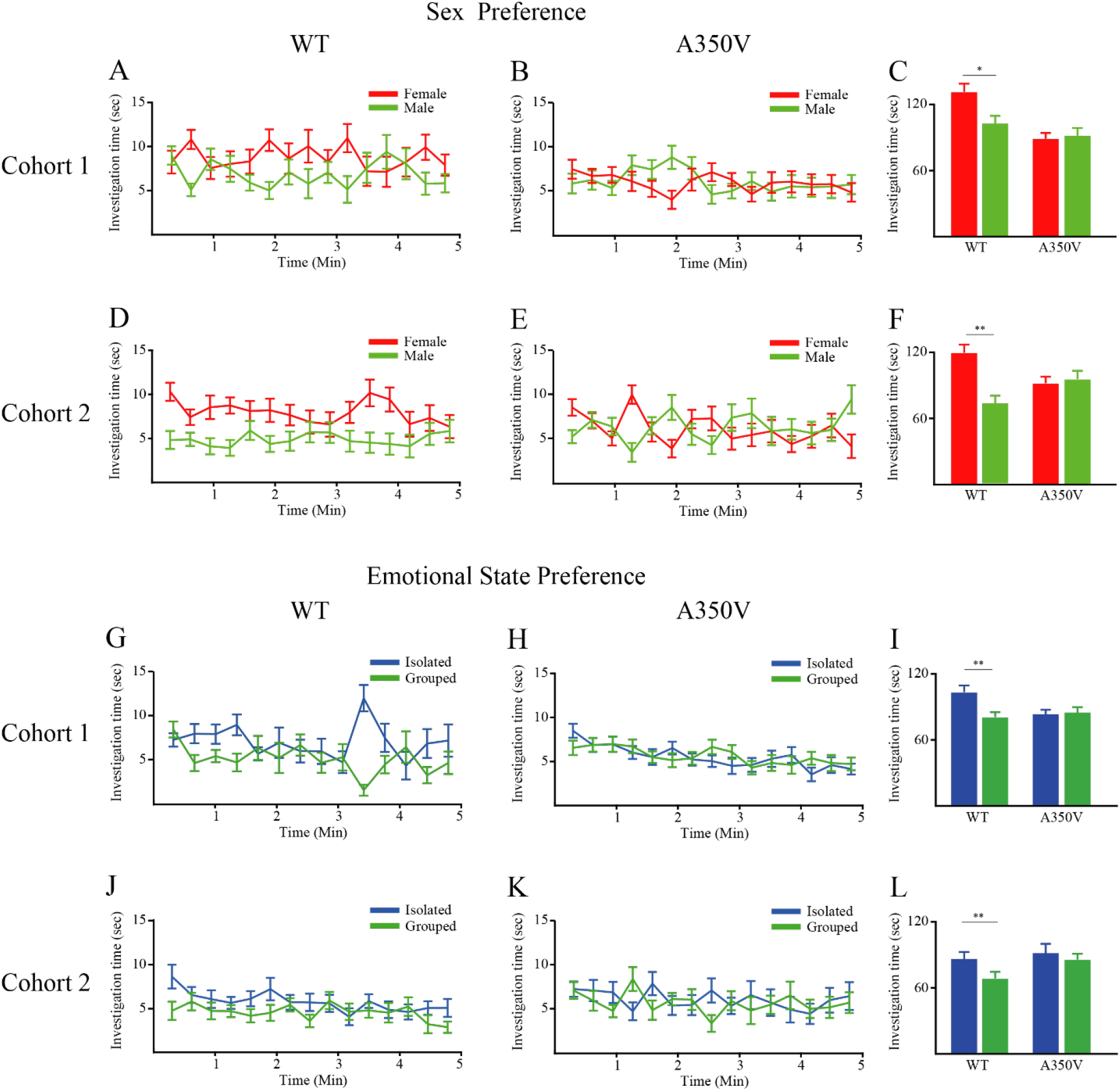
Deficits in sex preference and emotional state preference behaviors are replicated in two cohorts of A350V *Iqsec2* mice. (A-B) Mean investigation time measured separately for each stimulus (20-s bins) across the SxP session for the first cohort of WT (A, n=14) and A350V (B, n=20) mice. (C) Mean investigation time summed separately for each stimulus throughout the SxP test for WT (left) and A350V (right) subjects. *Post hoc* paired t-tests following main effect in mixed model ANOVA test. WT: t_13_=2.106, *p<0.05; A350V: t_19_=-0.274, p=0.787. (D-F) Same as A-C, for a second cohort of animals (WT. *post hoc* paired t-tests following main effect in mixed model ANOVA test. WT (n=19) t_18_=3.267; **p<0.01; A350V (n=16) t_15_=-0.307, p=0.763. (G-L) Same as A-F, for the ESP test. *post hoc* paired t-tests following main effect in mixed model ANOVA test. I, WT (n=11) t_10_=4.28; **p<0.01, A350V (n=28): t_27_=-0.172, p=0.865; L, WT (n=17) t_16_=3.11; **p<0.01, A350V (n=15) t_14_=-0.513, p=0.616.

**Figure S2.**
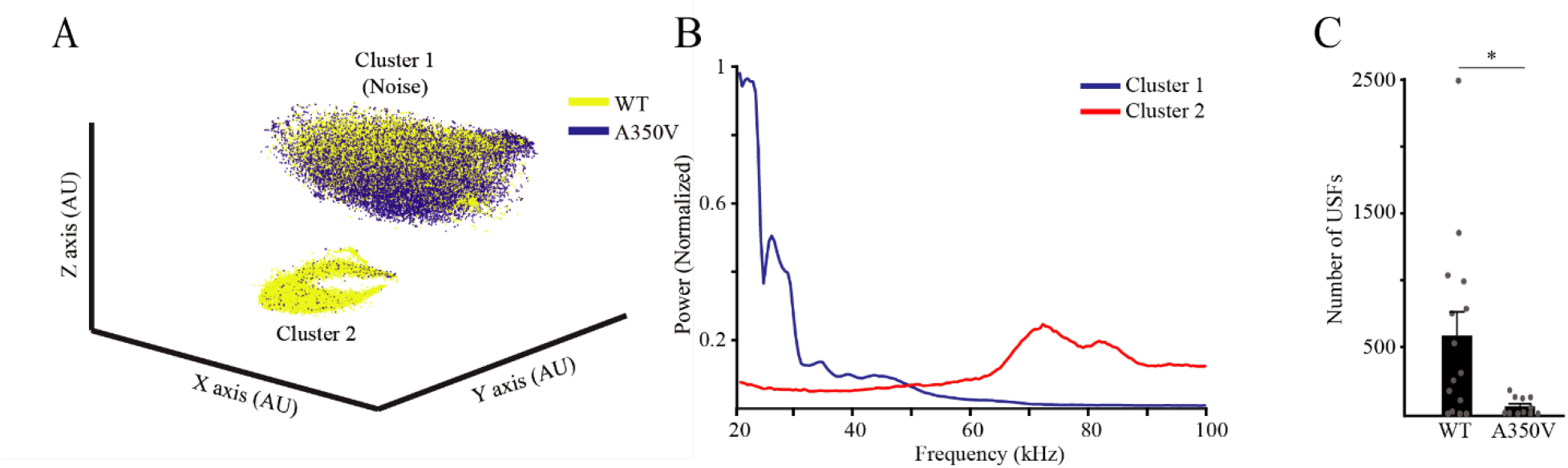
A350V *Iqsec2* male mice do not emit mating calls in the presence of a female mouse – A second cohort. (A) 3D t-SNE analysis of all ultrasonic fragments emitted by WT (yellow; n=15) and A350V (blue; n=11) adult male mice during an encounter with a female C57BL/6J mouse. Note that while cluster 1 (noise) seem to contain similar number of fragments from both genotypes, cluster 2 (vocalizations) contains almost only fragments from WT mice. (B) Power Spectral Density (PSD) analysis of all USFs recorded from all animals, calculated separately for each cluster. Note that in contrast to the PSD profile of cluster 1 which shows a wide range of frequencies, mainly at the lower range, the profile of cluster 2 shows a clear peak at the range of 60-90 kHz. (C) Comparison of the number of USFs from cluster 2 between the two genotypes. Note that A350V mice emitted significantly lower number of USFs as compared to WT animals (Mann-Whitney U test. U=39.000, *p<0.05).

**Figure S3.**
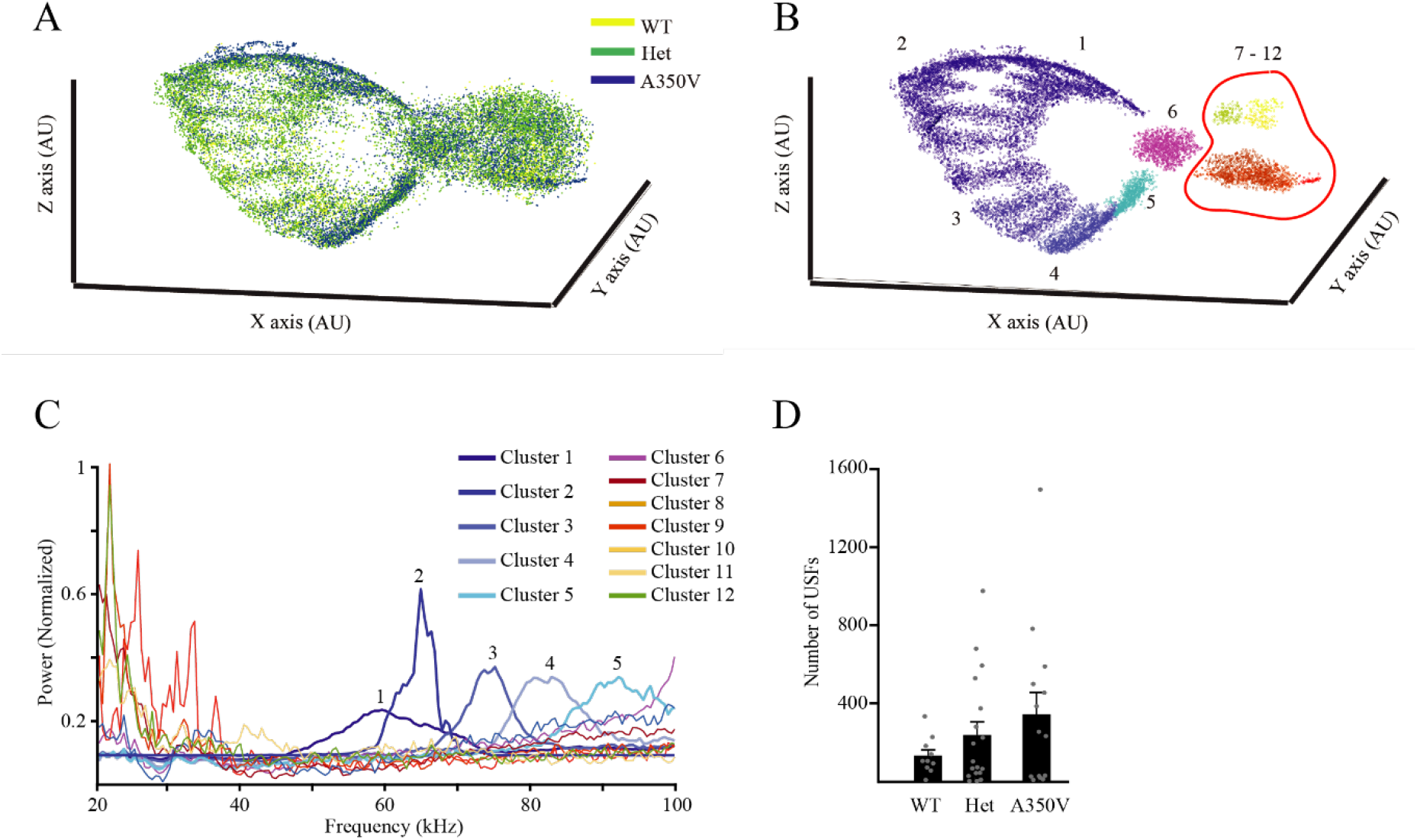
No difference in pup separation calls between WT and A350V *Iqsec2* littermates. (A) 3D t-SNE analysis of all ultrasonic fragments (USFs) recorded during pup separation experiments of WT (yellow; males and females; n=10), heterozygous (green; females; n=18) and A350V (blue; males; n=14) mice. All pups were genotyped after the recordings. (B) The same t-SNE analysis after automatic clustering (DBSCAN) of the various USFs. (C) PSD analysis of all USFs from each cluster for all animals analyzed in A. Note the while clusters 6-12 contain fragments of noise, clusters 1-5 contain fragments of genuine vocalizations with distinct peaks in the range of 50-100 kHz. (D) Comparison of the number of USFs from clusters 1-5 between the three groups of animals analyzed in A. There was no significant difference between the groups (Kruskal Wallis test, Chi=0.469, p=0.79).

**Figure S4.**
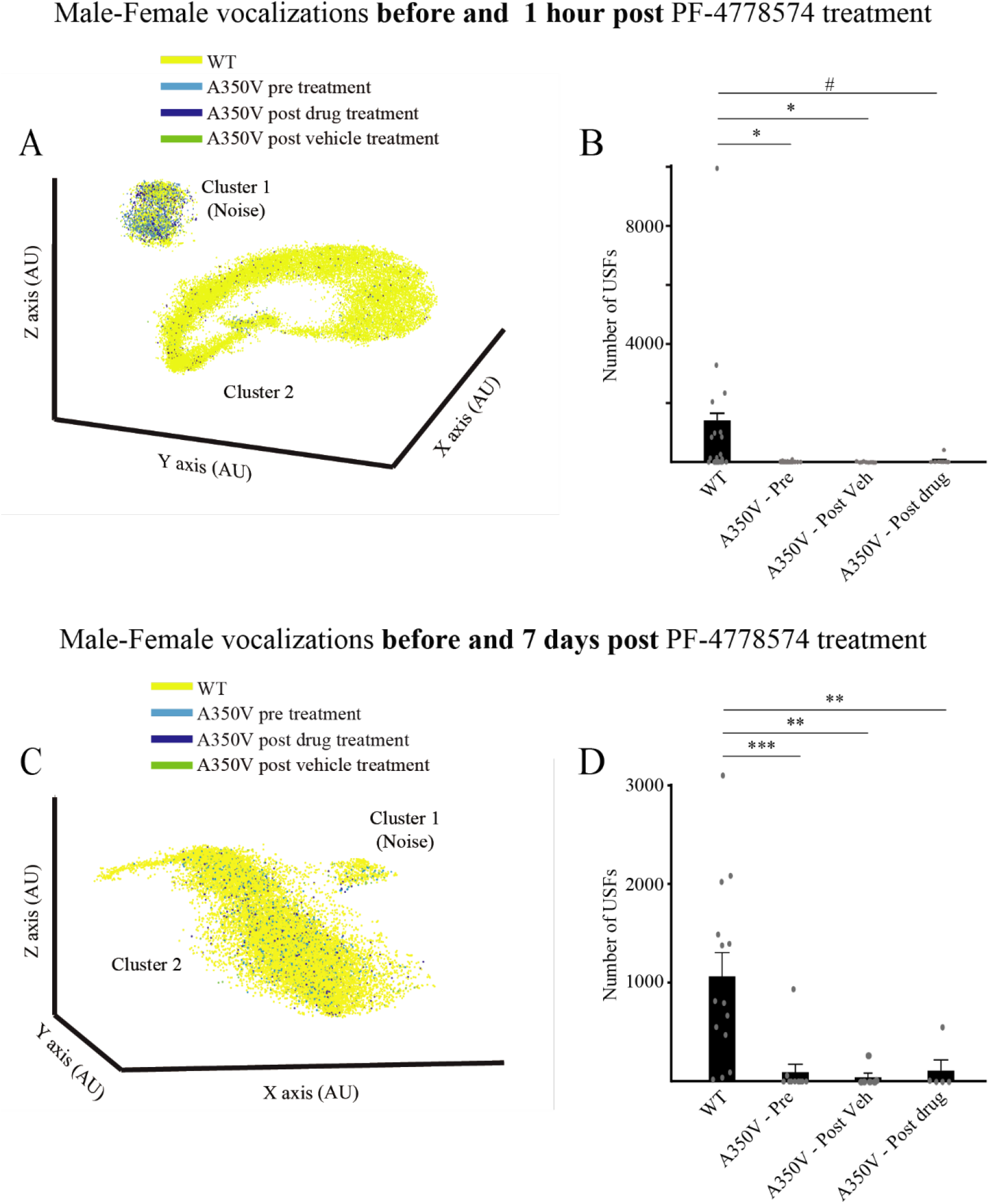
PF-4778574 does not affect the impaired mating calls in A350V *Iqsec2* mice. (A) 3D t-SNE analysis of all ultrasonic fragments (USFs) recorded during male-female interactions of WT mice (yellow; n=31), as well of A350V mice, before (light blue; n=36) or 1 hour after injection of either PF-4778574 (blue; n=11) or vehicle (green; n=10). Note that while cluster 1 (noise) seem to contain similar number of fragments from both genotypes, cluster 2 (vocalizations) contains almost only fragments from WT mice. (B) Comparison of the number of USFs from cluster 2 between the four groups of animals analyzed in A. Note that PF-4778574 administration did not elicit any effect on the number of USFs emitted by MT animals (Kruskal Wallis test, chi=8.737, p=0.033, post hoc Mann-Whitney U test following main effect: WT-MT before injection – U=390, p<0.05; WT-MT after PF-4778574 injection – U=81, p<0.05; WT-MT after vehicle injection – U=107.5, p=0.071). (C-D) As in A-B, for different cohorts of animals, with the vocalizations post treatments (PF-4778574 and vehicle injections) recorded 7 days following injections. Here too, no effect of PF-4778574 was observed. (WT - n=14, MT before injection - n=11, MT after vehicle injection - n=6, MT after PF-4778574 injection - n=5; Kruskal Wallis test, chi=20.31, p<0.001, *post hoc* Mann-Whitney U test following main effect: WT-MT before injection – U=10, p<0.001; WT-MT after PF-4778574 injection – U=6, p<0.01; WT-MT after vehicle injection – U=4, p<0.01).

## Notes

### Competing Interest Statement

The authors have declared no competing interest.

